# Acidosis, Zinc and HMGB1 in Sepsis: A Common Connection Involving Sialoglycan Recognition

**DOI:** 10.1101/2020.07.15.198010

**Authors:** Shoib S. Siddiqui, Chirag Dhar, Venkatasubramaniam Sundaramurthy, Aniruddha Sasmal, Hai Yu, Esther Bandala-Sanchez, Miaomiao Li, Xiaoxiao Zhang, Xi Chen, Leonard C. Harrison, Ding Xu, Ajit Varki

**Affiliations:** Departments of Medicine and Cellular and Molecular Medicine, University of California, San Diego, CA, USA; Glycobiology Research and Training Center, University of California, San Diego, CA, USA; Department of Chemistry, University of California, Davis, CA, 95616, USA; The Walter and Eliza Hall Institute of Medical Research, Parkville, Victoria 3052, Australia; Department of Medical Biology, University of Melbourne, Parkville, Victoria 3010, Australia; Department of Oral Biology, School of Dental Medicine, University at Buffalo, The State University of New York, USA

## Abstract

Blood pH is tightly regulated between 7.35-7.45, with values below 7.3 during sepsis being associated with lactic acidosis, low serum zinc, and release of proinflammatory HMGB1 from activated and/or necrotic cells. Using an *ex vivo* whole blood system to model lactic acidosis, we show that while HMGB1 does not engage leukocyte receptors at physiological pH, lowering pH with lactic acid facilitates binding. At normal pH, micromolar zinc supports plasma sialoglycoprotein binding by HMGB1, which is markedly reduced when pH is adjusted with lactic acid to sepsis levels. Glycan array studies confirmed zinc and pH-dependent HMGB1 binding to sialoglycans typical of plasma glycoproteins. Thus, proinflammatory effects of HMGB1 are suppressed via plasma sialoglycoproteins until drops in pH and zinc release HMGB1 to trigger downstream immune activation.

**Significance Statement:** HMGB1 sequestered by plasma sialoglycoproteins at physiological pH is released when pH and zinc concentrations fall in sepsis.

## Introduction

The pH of body fluids in healthy individuals spans a very broad range in different tissue types and organs, ranging from pH 1.5 (stomach contents), to 8.0 (urine). Human cells in tissue culture can also tolerate a wide range of pH values. In contrast, blood pH is tightly regulated between 7.35-7.45 (1), and departure out of this range (acidosis or alkalosis) can be very detrimental. For example, in the recent COVID-19 pandemic, 30% of non-survivors had acidosis, compared to 1% among survivors (2). Acidosis in sepsis is partly due to lactic acid release from anoxic tissues, which overwhelms the buffering capacity of circulating blood (3). A “cytokine storm “of proinflammatory mediators in sepsis triggers a cascade of destructive outcomes such as multiple organ failure (4–8) as currently seen in severe cases of COVID-19 infection (9). The mechanisms underlying lethality associated with low blood pH are not clear, but include low zinc levels and release from apoptotic or necrotic cells of HMGB1, a damage-associated molecular pattern (DAMP) defined as one of the late mediators of sepsis, further upregulating many other proinflammatory cytokines (10–12). Importantly, a recent study indicates HMGB1 levels are strongly associated with mortality in patients infected with SARS-CoV-2 (13). Here we show that sialylated plasma glycoproteins bind HMGB1 to suppress its ability to promote inflammatory responses in a zinc and pH-dependent manner. This finding provides an avenue for developing a new therapeutic strategy for treating sepsis.

## Results

### Mimicking lactic acidosis *ex vivo* in hirudin-anticoagulated whole blood

*In vivo* studies of acidosis and sepsis involve many complex factors and interactions. On the other hand, *ex vivo* reconstitution of purified blood components can result in artifacts, e.g., neutrophils get activated when separated away from erythrocytes and plasma (14). To study the significance of tightly regulated blood pH *ex vivo*, we sought to create a whole blood system mimicking lactic acidosis. Conventional anticoagulation with EDTA or citrate abrogates divalent cation functions, and heparin has many biological effects independent of anticoagulation. We have previously shown that the leech protein hirudin can be used to obtain whole blood anticoagulation *in vitro* (15). When lactic acid was added to freshly collected hirudin-anticoagulated whole blood, the pH first rose until a concentration of about 1 mM lactic acid was reached. Further addition then caused a sharp drop in blood pH. Such an initial rise in blood pH followed by a subsequent drop is seen in patients with sepsis (16). To further develop this model, we introduced HMGB1, a DAMP (17–19) associated with poor prognosis in late sepsis (20, 21).

### Neutrophils in whole blood are activated by HMGB1 at low pH due to better binding, and activation is attenuated with an HMGB1 blocking-antibody

Interaction of HMGB1 with Toll-like receptors (TLRs) during sepsis is well documented (22). The proinflammatory activity of HMGB1 is due to binding to targets such as TLR-2, TLR-4, TLR-9 and RAGE that are expressed on leukocytes and endothelial cells (23, 24). We, therefore, introduced exogenous HMGB1 into our whole blood acidosis model and tracked CD11b expression on neutrophils, as a sensitive marker of activation triggered by HMGB1. Increased neutrophil activation was noted when HMGB1 was incubated with whole blood at low pH as compared to physiological pH (Figure 1A). This effect was partially attenuated by adding HMGB1 blocking antibody (Figure 1B). Enhanced activation at low pH coincides with increased HMGB1 binding to neutrophils and monocytes (compare upper and lower panels of Figure 2A and B). Thus, physiological blood pH limits interaction of HMGB1 with leukocyte receptors, suggesting natural inhibitor(s) of HMGB1 interaction in blood. Looking for candidate inhibitors, we noted earlier evidence that HMGB1 can interact with CD24 and CD52, two heavily sialylated proteins (25, 26) in a trimolecular complex with Siglec-10, a known sialic acid-binding protein. CD52-Fc bound specifically to the proinflammatory Box B domain of HMGB1, and this, in turn, promoted binding of the CD52 N-linked glycan sialic acid with Siglec-10 (26). Furthermore, sialidase treatment abolished CD52 binding to HMGB1, indicating that it might be a sialic acid-binding lectin. Since normal blood plasma contains ∼2 mM sialic acid attached to glycans on plasma proteins, (27), we hypothesized that the unknown natural inhibitor might be the sialome (the total sum of all sialic acids presented on plasma glycoproteins).

**Fig 1:**
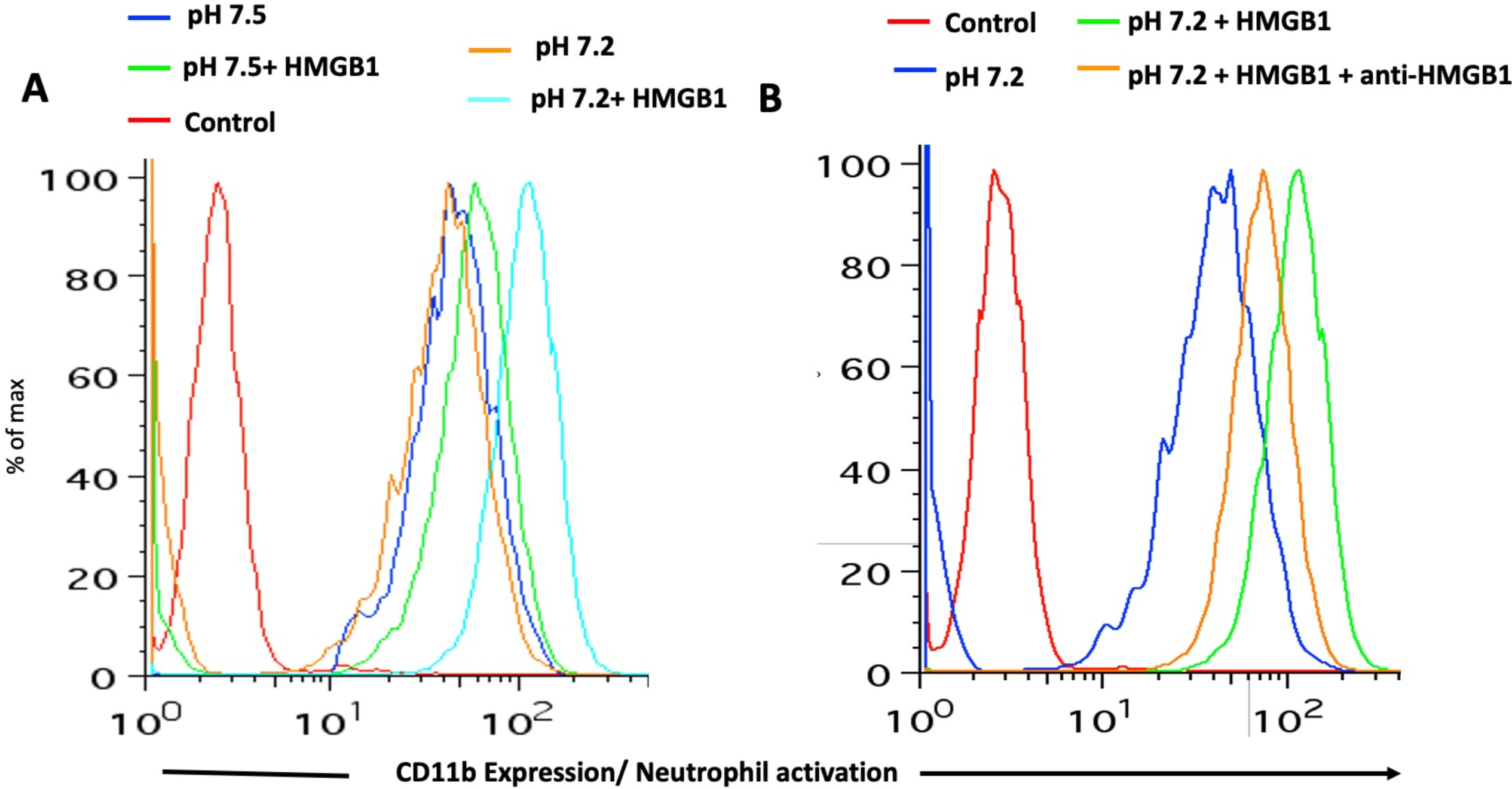
Mimicking sepsis by adding lactic acid to whole blood triggers activation of neutrophils by HMGB1, which is partially attenuated by an HMGB1-blocking antibody: CD11b expression was determined by flow cytometry after incubating whole blood with/without HMGB1 (1 µg/ml). A) Neutrophils are activated when incubated with HMGB1 in whole blood at pH 7.2 (Chromatograms: Red-Control, Blue-whole blood at pH 7.5, Orange-whole blood at pH 7.2, Green-whole blood at pH 7.5 with HMGB1, Cyan-whole blood at pH 7.2 with HMGB1). B) Activation is partially attenuated with an HMGB1-blocking antibody (50 µg/ml) (Chromatograms: Red-Isotype control, Blue-whole blood at pH 7.2, Green-whole blood at pH 7.2 with HMGB1, Orange-whole blood at pH 7.2 with HMGB1 and an HMGB1-block

**Fig 2:**
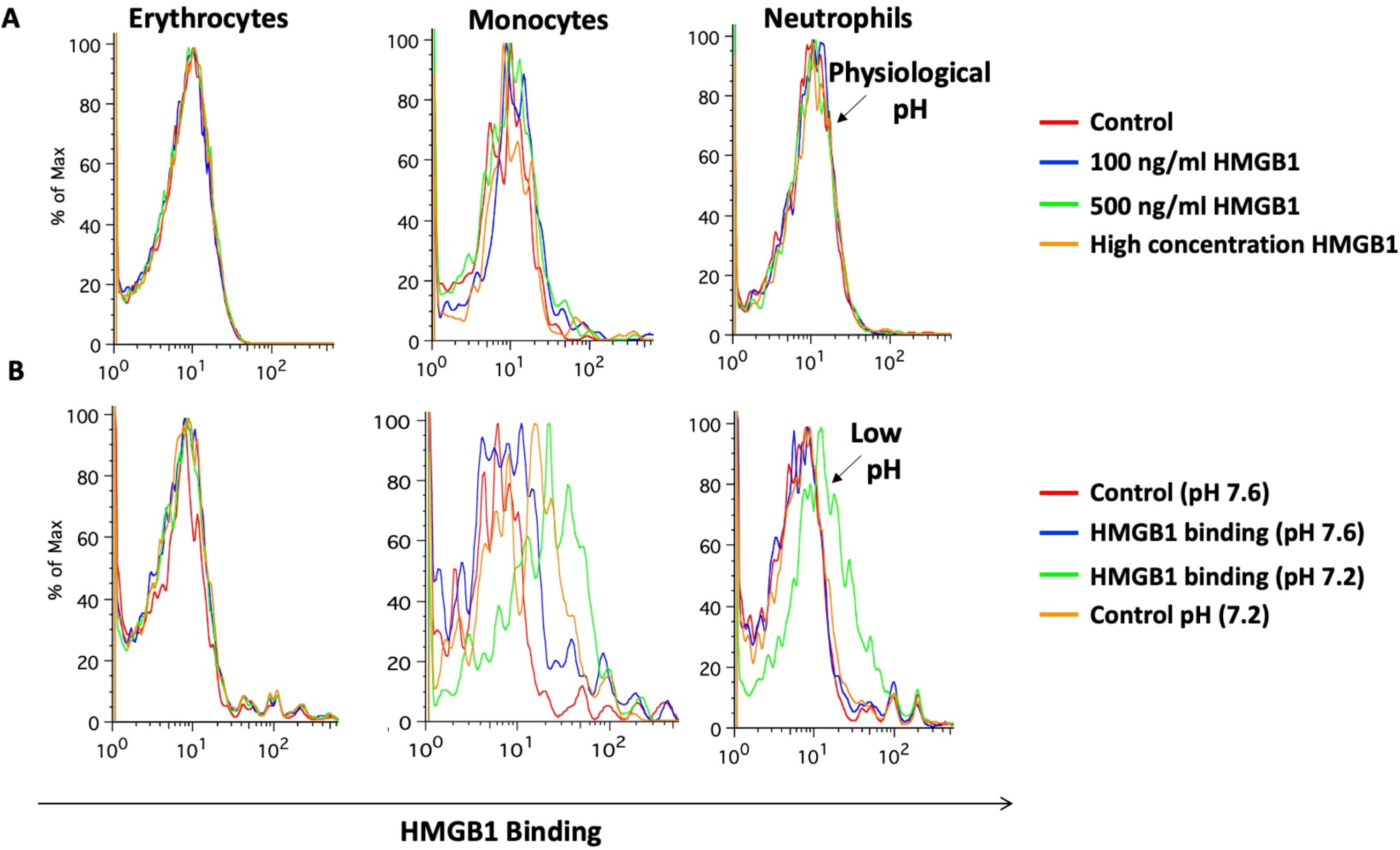
Mimicking sepsis by adding lactic acid to whole blood triggers binding of HMGB1 to leukocytes: A) Ability of HMGB1 to bind to different cell types of the blood (erythrocytes, monocytes and neutrophils) was determined by using different concentrations (100 ng/ml, 500 ng/ml and 5 µg/ml) of HMGB1 at physiological conditions. B) Different cell types of blood were used for binding with HMGB1 (100 ng/ml) at physiological and lower pH (pH 7.2, adjusted with lactic acid).

### Among divalent cations, only zinc supported the robust binding of HMGB1 with sialylated glycoproteins at physiological pH

The binding buffer used in prior HMGB1 studies included millimolar concentrations of manganese cation (Mn^2+^), a feature likely carried over from the unrelated function of nuclear HMGB1 binding to DNA. Looking at earlier studies of the interaction of HMGB1 with CD24 and CD52, we noticed that all those experiments were performed in a buffer containing millimolar Mn^2+^ concentrations (25, 28–30). These concentrations were very high in comparison with the physiological levels of Mn^2+^ in the blood (4-15 µg/L). We predicted that there might be other divalent cation(s) that are better co-factor(s) for HMGB1 and facilitate its binding with sialic acids. Indeed, upon testing micromolar concentrations of many divalent cations, we found that only zinc cation (Zn^2+^) supported robust binding with sialylated glycoproteins (Figure 3A). We tested α1-acid glycoprotein and 3’-sialyllactose as binding partners for HMGB1 in the presence of different cations and again found that only Zn^2+^ facilitated binding. There was a modest binding of 3’-sialyllactose with HMGB1 in the presence of Mn^2+^, but robust binding was only seen with Zn^2+^-containing buffer (Figure 3B).

**Fig 3:**
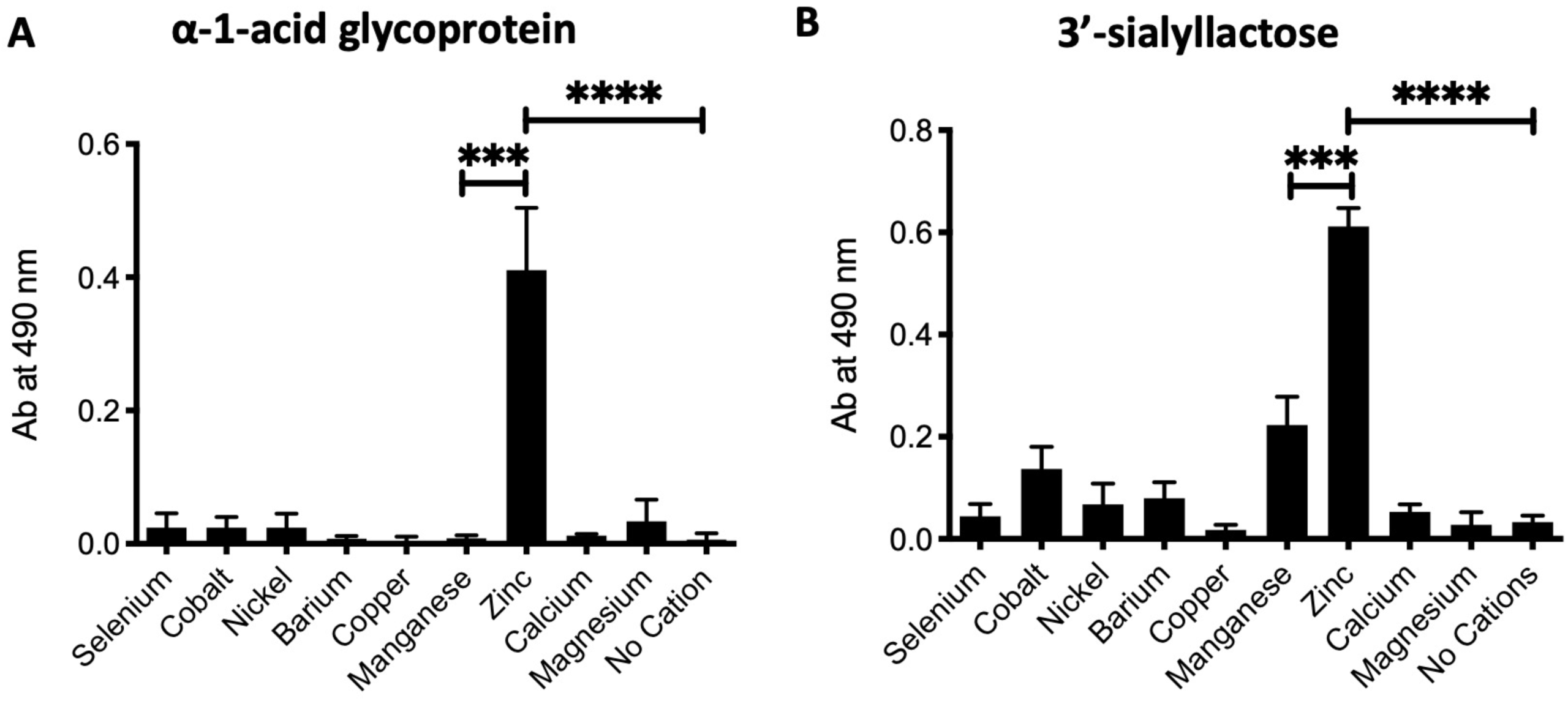
Among divalent cations, only zinc supported robust binding of HMGB1 with sialylated glycoproteins: A, B) Many divalent cations were used in binding buffer with concentration 500 µM and binding with human α1-acid glycoprotein and 3’-sialyllactose was determined using ELISA. The experiments were performed in triplicates where data shows mean±SD.

### Replacing plasma with buffer at physiological pH allows HMGB1 to activate neutrophils, suggesting sequestration by plasma sialoglycoproteins

We next asked which whole blood components were preventing neutrophil activation under physiological conditions. Hirudin-anticoagulated whole blood at physiological pH was spun down and plasma either replaced with HEPES buffer (pH 7.5) supplemented with Zn^2+^ or with the same plasma that had been removed. After incubating with HMGB1, neutrophils were in a more activated state when incubated in the buffer as compared to when plasma was added back (Figure 4A). Independent studies have shown that HMGB1 binds to sialic acid on glycoproteins (26, 31) and we posited that the ∼2 mM bound sialic acid present on plasma glycoproteins might lead to sequestration of HMGB1 under physiological condition. We also tested the effect of pH on the binding of HMGB1 to α1-acid glycoprotein and found that optimal binding was at physiological pH, with less binding at pH 7.2 with buffer containing Zn^2+^ (Figure 4B).

**Fig 4:**
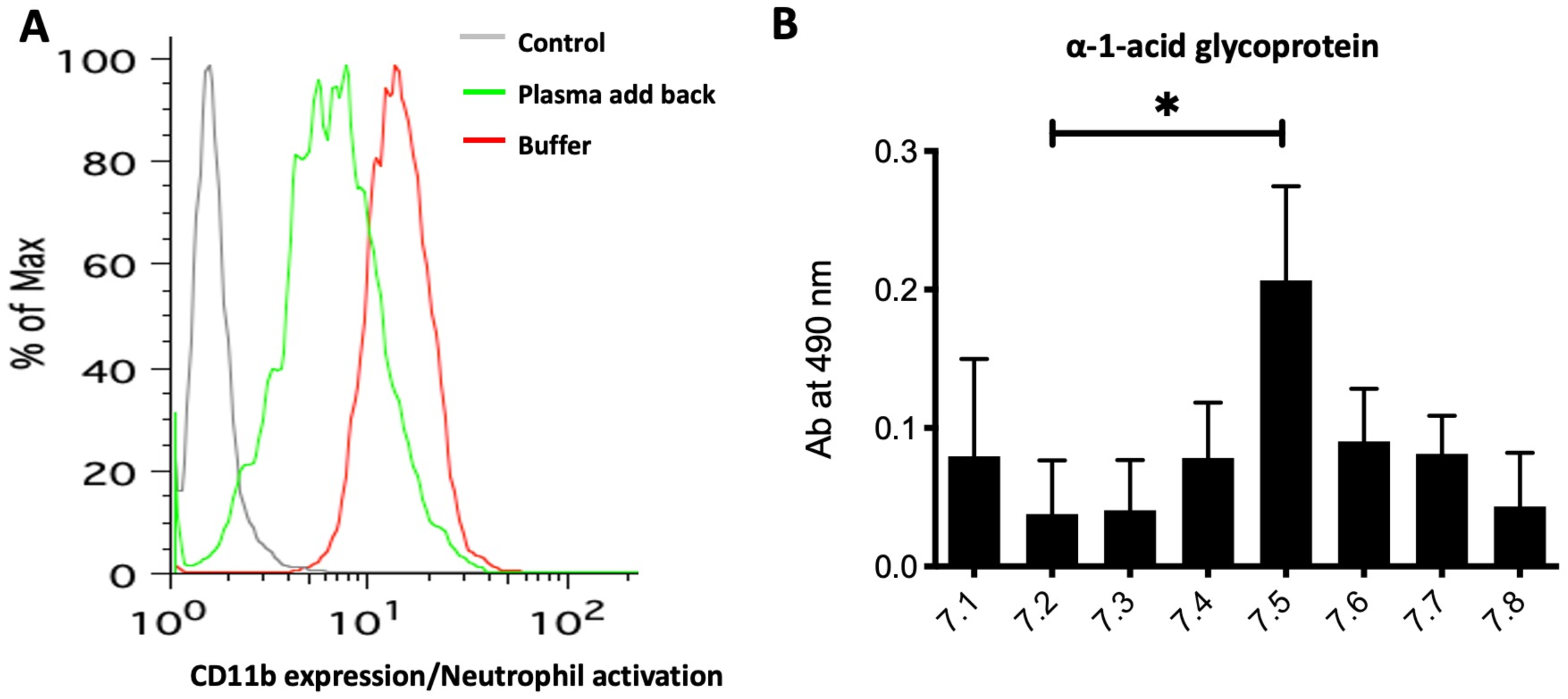
Replacing plasma with a buffer at physiological pH allows HMGB1 to activate neutrophils. A) 1 ml of blood was drawn from a healthy individual and spun down. The plasma was replaced with HEPES buffer containing zinc (500 µM of Zn^2+^) or plasma was added back. The CD11b expression as a marker of neutrophil activation was measured. B) The binding of HMGB1 to α1-acid glycoprotein was checked with a binding buffer using different pH ranging from 7.1 to 7.8.

### Sialoglycan array studies of HMGB1 confirm that it is a sialic acid-binding lectin with optimal binding at physiological blood pH in the presence of zinc cations

We previously reported a sialoglycan microarray platform used to identify, characterize, and validate the Sia-binding properties of proteins, lectins, and antibodies (32–34). After identifying Zn^2+^-dependent HMGB1 binding to sialoglycoproteins, we next investigated the ability of HMGB1 to bind with multiple sialoglycans abundantly found in plasma proteins. We performed sialoglycan array studies of HMGB1 under four different conditions: 1) At physiological pH with Zn^2+^, 2) At physiological pH without Zn^2+^, 3) At pH 7.2 with Zn^2+^ 4) At pH 7.2 without Zn^2+^. These array studies further confirmed the binding of HMGB1 with multiple sialylated glycan sequences that are typically found on plasma glycoproteins, in pH- and Zn^2+^-dependent fashion (Figure 5A and 5B respectively). Additionally, we checked the binding of HMGB1 to sialic acids in sialoglycan microarray using 0, 15 and 150 µM concentrations of Zn^2+^ and observed a dose-dependent effect (Figure 5B). This assay showed the relevance of Zn^2+^ in this binding phenomenon at a physiological concentration (∼15 µM). On resolving the binding of HMGB1 at physiological pH and in the presence of zinc, the binding on the microarray was exclusively to sialylated glycans confirming our findings (Figure 5C). A heat map representation of all these findings and HMGB1 binding to individual glycosides is provided in supplementary figures 3 and 4 respectively.

**Fig 5:**
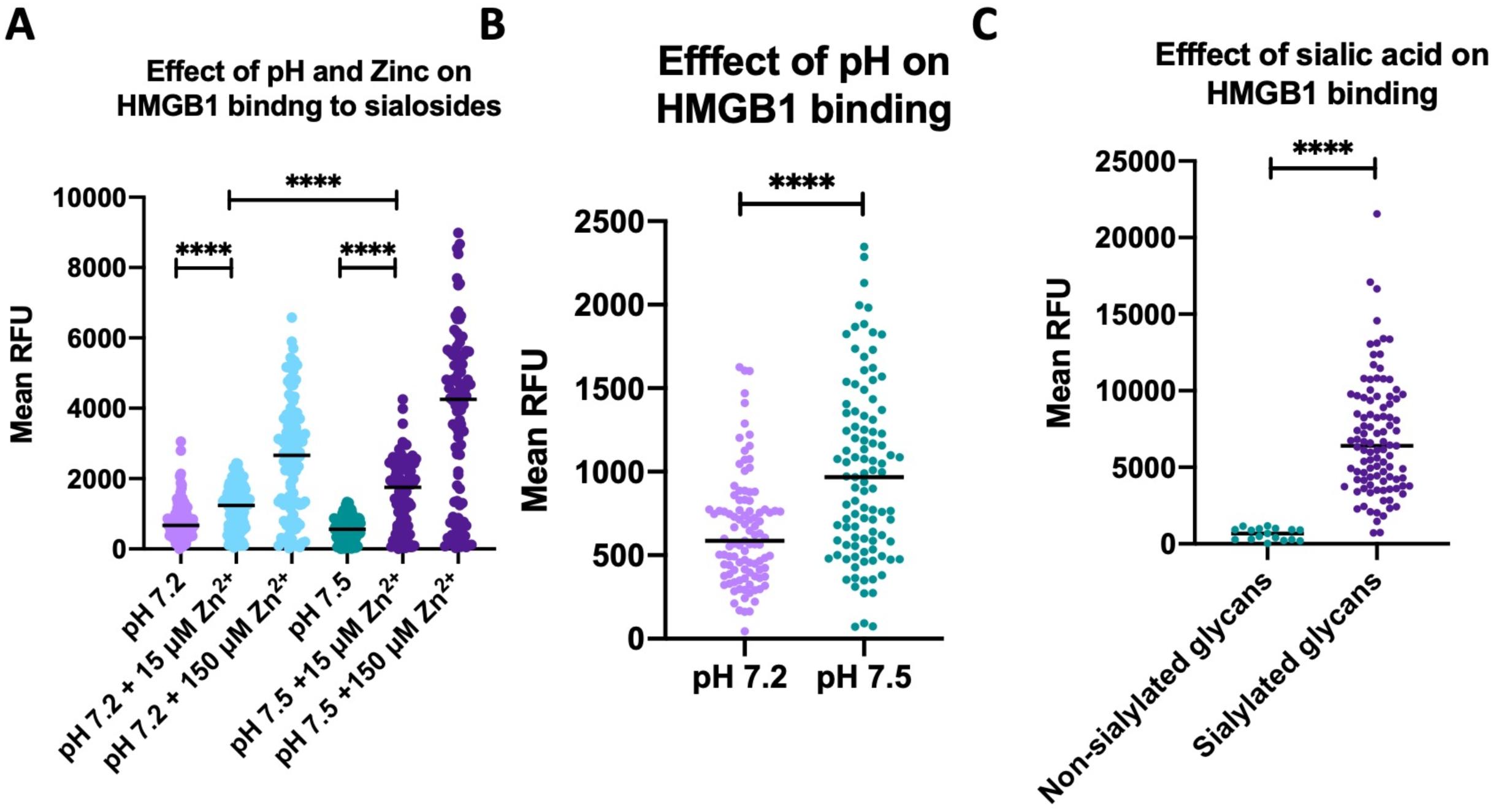
Sialoglycan array studies of HMGB1 confirm that it is a sialic acid-binding lectin with optimal binding at physiological blood pH in the presence of zinc: A) The sialoglycan array was performed to test the binding of HMGB1 with multiple sialylated probes. The binding buffer used for the assay either contained zinc and pH 7.5, no zinc, pH 7.5, with zinc, pH 7.2 and no zinc at pH 7.2. The concentration of zinc used was 15 µM and 150 µM. (Representative image of a single experiment. Wilcoxon matched pairs signed rank test used. **** represents p-value < 0.0001) B) Additional microarray experiments with 500 µM further resolve the pH-dependent binding difference. (Representative image of the mean of two experiments. Unpaired t-test with Welch’s correction used to compare the two groups. **** represents p-value < 0.0001) C) The difference of HMGB1 to sialosides and non-sialosides at physiological pH in the presence of 500 µM zinc. (Representative image of a single experiment where zinc was used at a concentration of 500 µM. Kolmogorov-Smirnov test used. **** represents p-value < 0.0001)

### Heparin, a previously known anionic glycan binding partner of HMGB1, does not exhibit pH sensitivity, and Zn^2+^ only partially facilitates binding

HMGB1 is known to bind heparin, a heavily sulfated glycan carrying many negatively charged groups (35, 36). We checked the binding of HMGB1 with heparin at different pH values and found that unlike binding with Sia, it was not pH-sensitive (Supplementary Figure 1A). Moreover, there was appreciable baseline binding of HMGB1 with heparin that only increased partially with Zn^2+^ supplementation (Supplementary Figure 1B). This data indicates that the binding of heparin and sialic acid are very different. The B-Box of HMGB1 that mediates sialic acid binding (26) has three arginine residues (26) that might be involved in sialic acid recognition. We made single mutants of arginine residues at positions 97, 110 and 163. When we checked the sialic acid binding, we could not find any difference between either of the mutants and WT HMGB1 (Supplementary Fig 2). We suspect other positively charged residues and/or multiple arginines to mediate sialic acid binding.

## Discussion

Here we report one plausible explanation for the tight regulation of blood pH between 7.35-7.45, showing that even a slight reduction to pH 7.2 abolishes the zinc-dependent sequestration of HMGB1 by plasma sialoglycoproteins, releasing it to bind to activating receptors on neutrophils. HMGB1 was originally discovered in the cell nucleus (37–40), playing a role in DNA bending, replication and transcription (41, 42). Much later, HMGB1 was found to be passively or actively released in conditions like sepsis, leading to inflammation (21, 41, 43). i.e, it is as a DAMP (44). HMGB1 retention inside the nucleus is dictated by conserved lysine residues (45). Inflammatory stimuli trigger acetylation of these lysine residues and trafficking of HMGB1 to the cytosol, and eventually to the extracellular space. The different domains of HMGB1 are Box A, Box B and an acidic tail. While Box A and Box B possess many arginine and lysine residues, the acidic tail is enriched with glutamic and aspartic acid residues. Box B is proinflammatory whereas Box A behaves like an antagonist and mimics an anti-HMGB1 antibody (26, 46).

While TNF-α and IL-1β are released early during sepsis, HMGB1 is a late mediator expressed only after about 24 hours and remains at elevated levels before death occurs (47). Many preclinical studies show protection against sepsis upon injection of blocking antibodies of HMGB1 or just injection of Box A protein (48). The proinflammatory activity of HMGB1 is well studied. However, the anti-inflammatory activity of HMGB1 also has been documented in multiple studies (49–51). Recently, it was shown that HMGB1 binds soluble CD52 and this complex binds with Siglec-10 on T-cells leading to SHP-1 (phosphatase) recruitment that dephosphorylates LCK and Zap70, thus activating an anti-inflammatory cascade (26, 52). In addition, haptoglobin (49), C1q and TIM3 also show anti-inflammatory activity of HMGB1 (50, 51).

In this study, we found that at physiological blood pH, there is no interaction of HMGB1 with its receptors on leukocytes. Surprisingly, when we lowered the pH using lactic acid (to mimic lactic acidosis, a characteristic feature of sepsis), the interaction was restored. Furthermore, the high concentration of sialic acids in plasma glycoproteins was found to be the likely inhibitor of interactions between HMGB1 and TLRs. We further characterized the role of HMGB1 as a sialic acid-binding lectin and found that zinc is a required co-factor. Moreover, we confirmed all our findings with lipopolysaccharide (LPS)-free HMGB1 and used a glycan array that detected the binding of HMGB1 with several sialic acid probes (See Supplemental Table 1) in a pH and zinc-dependent manner.

Taken together, our findings lead us to propose that under physiological conditions (pH 7.35-7.45) and normal zinc concentrations, there is a potent binding of HMGB1 with plasma sialoglycoproteins (Figure 6A). Under septic conditions, drops in pH and zinc concentration decrease interactions between HMGB1 and plasma sialoglycoproteins leading to the liberation of HMGB1 to bind with TLRs, to enhance inflammation (Figure 6B). Therefore, proinflammatory and anti-inflammatory activities of HMGB1 are the two sides of the same coin and are dependent on the different physiological conditions. While the proinflammatory role of HMGB1 is very well studied, recent studies have reported an anti-inflammatory role for HMGB1 (25, 50–52). The exact mechanism that enables HMGB1 to switch from its proinflammatory to anti-inflammatory role and vice-versa is not very well described. One factor known to enable its switch from being proinflammatory to anti-inflammatory is its oxidative state. The disulfide form of HMGB1 is proinflammatory, and the sulfonate form is involved in the resolution of inflammation (53–55). In the current study, we have identified another mechanism by which HMGB1 switches from its proinflammatory to anti-inflammatory role in a pH- and zinc-dependent manner. Sepsis is characterized by a decrease in pH and zinc concentration of the blood. We hypothesize that under physiological conditions, HMGB1 binds with sialoglycoproteins of blood keeping it in a quiescent state. During sepsis, the drop in pH and zinc concentration of the blood leads to disruption of HMGB1’s binding with sialic acid, enabling the free HMGB1 to bind with TLRs and RAGE present on immune cells and the endothelium. This activates a cascade of the inflammatory response, which if untreated, might lead to multiple organ failure or even death.

**Fig 6:**
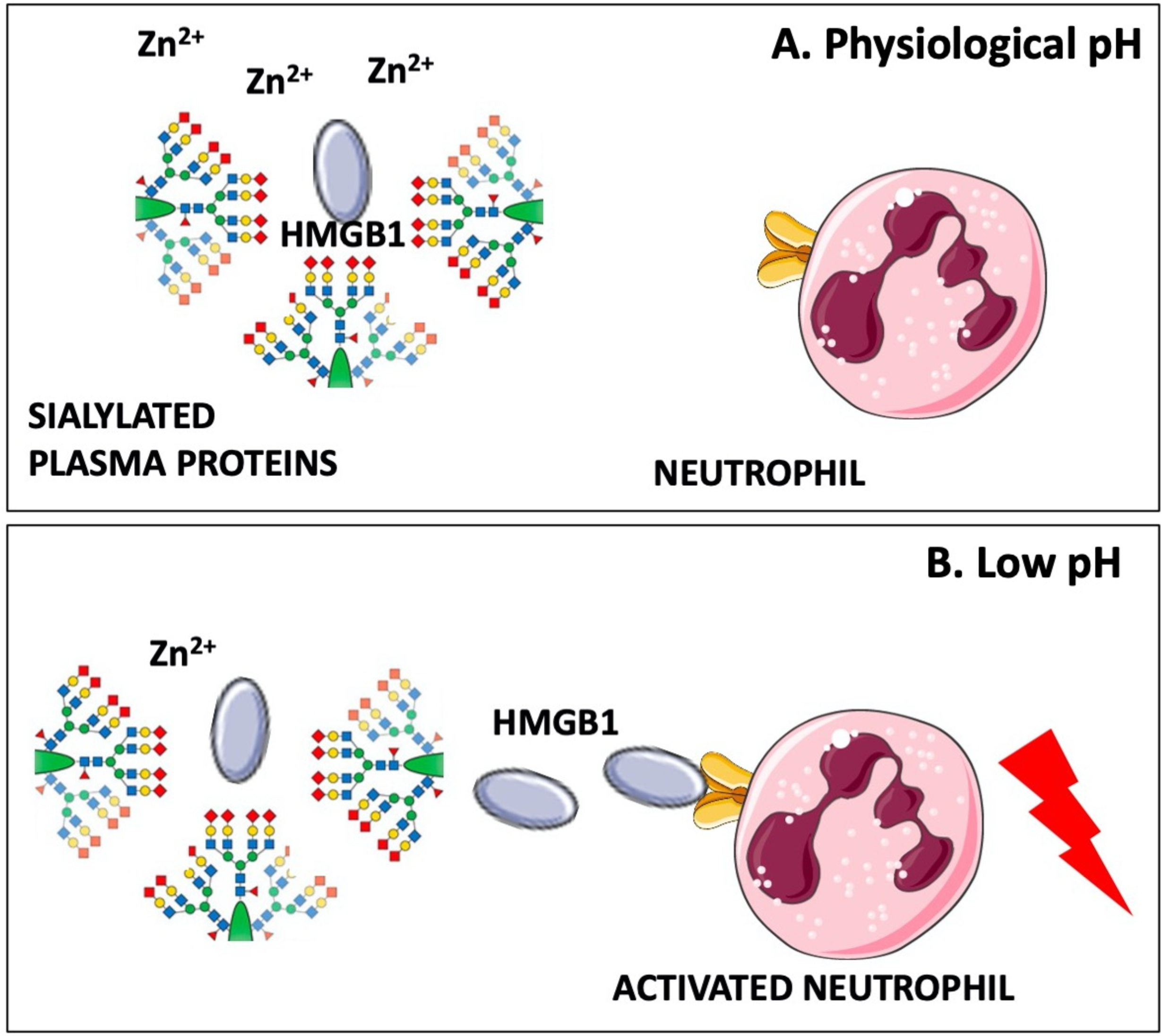
Proposed model of sequestration of HMGB1 by sialoglycoproteins to prevent HMGB1 binding to receptors on leukocytes: A, B) A schematic showing the binding of HMGB1 to sialic acid under physiological pH and binding to leukocyte receptors at low pH.

Also consistent with our hypothesis are the findings that survival in mouse models of sepsis can be improved by infusion of soluble CD52(56), and that the sialic acid binding feature of HMGB1 is restricted to the disulfide-form of HMGB1 (26), which is expected to be formed when the cytosolic reduced form is released into the oxidizing environment of the bloodstream. We suggest that the potent proinflammatory effects of HMGB1 are normally kept in check via sequestration by plasma sialoglycoproteins at physiological pH and zinc levels and is triggered when pH and zinc levels fall in the late stages of sepsis. In this regard, it is notable that the acute phase response to inflammation results in high production of hypersialylated molecules such as α1-acid glycoprotein from the liver and endothelium, which may then act as a negative feedback loop (57–59). Current clinical trials that are independently studying zinc supplementation (ClinicalTrials.gov Identifier: NCT01328509 NCT02130388) or pH normalization (NCT03530046) may be more successful if these approaches are combined, and perhaps supplemented by infusions of heavily sialylated molecules like CD52. Additionally, studies evaluating plasma exchange in subjects with septic shock (example NCT03366220) may show superior efficacy if supplemented with zinc infusions and pH correction. Pre-clinical studies are presently evaluating a function blocking anti-HMGB1 antibody (60). We performed our assays with HMGB1 purchased from HMG biotech, also produced it in *E. coli* and finally confirmed findings using HMGB1 expressed in 293 Freestyle cells. In order to recapitulate the characteristics of HMGB1 in septic conditions, we used the disulfide linked form in all our assays. Future studies should address whether other post-translational modifications such as acetylation, methylation, phosphorylation or oxidation have any further effect on HMGB1’s propensity to bind sialic acids.

Numerous studies have shown that zinc is protective against sepsis (61–63). Additionally, blood zinc levels usually decrease during inflammation because it is sequestered to the nucleus where it is required as a co-factor for expression of proinflammatory genes and proteins (61, 64, 65). Thus, lowering of the zinc level in blood is detrimental. The mechanism of action for the anti-inflammatory effect of zinc is extensively studied. These include effect impact on the microbiome, lowering of NF-κB levels, chemotaxis and phagocytosis by immune cells, anti-oxidative stress and adaptive immune response (61).

In this regard, it is notable that a recent study also shows the role of zinc, pH and ionic strength on the oligomerization of HMGB1 (66). We did not investigate any role of zinc or pH on the structural changes or oligomerization of HMGB1. It seems that at particular pH and zinc concentration, a positively charged residue of HMGB1 is exposed for binding with sialic acid. This residue may not be surface available at lower pH and low zinc concentration. In this study, we could not pinpoint the critical residue that is important for sialic acid binding.

HMGB1 has been reported to bind many ligands and some of which are highly negatively charged molecules such as heparin/heparan sulfate (35). We wanted to determine if the interaction of HMGB1 with sialic acid, which is also negatively charged, is a generic electrostatic charge-based interaction. Therefore, we tested the binding of HMGB1 with the acidic glycosaminoglycan, hyaluronan, but could not detect any binding (data not shown). Upon testing with heparin, we found that while HMGB1 did bind with heparin, it did not show any pH dependency. Moreover, binding was only partially enhanced in the presence of zinc. This shows that a different set of amino acid(s) might be required for binding to heparin and sialic acid. Notably, under physiological conditions, sialic acid is present in the blood, but the concentrations of other anionic glycans (heparan sulfate, hyaluronic acid etc.) are low.

Our findings, if confirmed in randomized clinical trials, have broad implications in the management of sepsis and possibly other types of acidosis. Sepsis is a significant cause of mortality, with a recent study implicating it as the cause of twice as many deaths as earlier estimated (67). These findings are of particular importance in light of the present COVID-19 pandemic/survivorship in these patients. Acute respiratory distress syndrome (ARDS), a deadly complication of the SARS-CoV-2 and SARS-CoV-1, has been linked with HMGB1 production (68–70). Recent articles suggest a potential link between HMGB1 and the pathogenesis of COVID-19 (71, 72). A recent study showed that HMGB1 strongly correlates with mortality in COVID-19 patients (13). Additionally, another recent study showed 100% of COVID-19 non-survivors had sepsis and 30% of these had acidosis (2). While the Surviving Sepsis Campaign does not suggest the use of convalescent plasma in critically ill patients, (73), the FDA has approved its use as an investigational new drug. A small study of five critically ill COVID-19 patients treated with convalescent plasma showed improvements in sepsis related SOFA scores (74). A clinicaltrials.gov search for “COVID “and “convalescent plasma “on April 6, 2020 yielded 9 results of trials ranging from phase 1 to phase 3. While the circulating antibodies are likely to be beneficial on their own, the HMGB1-sequestering properties of plasma sialoglycoproteins may also contribute to suppressing the “cytokine storm “. These effects are likely to be further enhanced if plasmapheresis is supplemented with aggressive pH correction and zinc supplementation.

## Material and Methods

### ELISA for binding of HMGB1 with α1-acid glycoprotein or 3’-sialyllactose

500 ng-1µg of HMGB1 recombinant protein (HMG Biotech) diluted with the binding buffer (20 mM HEPES, 150 mM NaCl, 500 µM ZnCl_2_) was immobilized by applying on a 96 well flat bottom plate (Corning co-star, Catalogue number 9018) and incubating overnight at 4 °C. The wells were washed thrice with 200 µl of binding buffer per well, followed by blocking with 150 µl of 5% BSA (prepared in binding buffer). The plate was then incubated at room temperature (RT) for 1 hour with shaking. The blocking solution was removed by flicking plate and tapping at a dry paper towel. Then 1 µg/well of biotinylated α_1_-acid glycoprotein (Sigma, Catalogue number-112150) or 3’-sialyllactose-PAA-biotinylated (Glycotech, Catalogue number-01-038), diluted in binding buffer, was applied on every well except the secondary antibody control wells which were left with only binding buffer. The plate was incubated 1-2 hours at RT on the shaker. The solution was removed, and wells were washed thrice with 200 µl binding buffer per well. The secondary antibody (Streptavidin-HRP, abcam, Catalogue number-ab7403-500) was applied at a dilution of 1:20000 in binding buffer and the plate was incubated for 1 hour at RT with shaking. Then O-phenylenediamine (OPD) based substrate solution for HRP was prepared by adding 5 mg of OPD and 25 µl of 30% H_2_O_2_ to 15 ml of Citrate-PO_4-_buffer. 140 µl of OPD substrate solution was added to each well and incubated in the dark until color development. Upon color development, the reaction was stopped using 40 µl of 2N H_2_SO_4_ and the absorbance was acquired at 490 nm with a plate reader. For the ELISA with different divalent cations, the binding buffer was prepared using the particular cation containing salt instead of ZnCl_2_. Each incubation and wash was performed using the respective binding buffer.

### Hirudin-anticoagulated whole blood assays

Informed consent was obtained from healthy individuals as per a protocol approved by the UCSD Human Research Protection Programs Institutional Review Board and venous blood was collected in hirudin coated tubes (ThermoFischer catalogue number-NC1054637). Hirudin was chosen as the anticoagulant as EDTA and heparin interferes with normal bioprocesses (chelation by EDTA and binding to and modulating cell-surface proteins by heparin). The pH of blood, when measured at the start of various assays varied between 7.5 and 7.6 and is referred to as the “physiological “pH.

### Flow cytometry analysis for HMGB1 activation of/binding to leukocytes

To test for neutrophill activation, 100 µl of whole blood was incubated with 1 µg/ml of HMGB1 for 30 minutes at 37 °C. CD11b expression was measured by flow cytometry as described earlier (14, 75). Blocking with an anti-HMGB1 antibody (Clone 3E8, BioLegend, catalogue number-651402) was performed with 50 µg/ml antibody as described earlier (60). For plasma addback studies, whole blood was spun down at 500 x g for 5 minutes and replaced with HEPES buffer supplemented with 500 µM ZnCl_2._ Binding assays were performed with 500 µl of whole blood. The required amount of HMGB1 (0, 100, 500, 5000 ng/ml) was added to 500 µl of blood and incubated at 37^°^ C for 60 minutes with rotation. After centrifuging at 600 x g for 5 minutes, the cells were washed with 1 ml of PBS and finally resuspended in 100 µl of FACS buffer (1% BSA in PBS with Ca^2+^/Mg^2+^) with anti-HMGB1 antibody (10µg/ml, BioLegend, catalogue number-651402). The cells were incubated at 4 ^°^C for 30 minutes on ice and were washed with 1 ml PBS (containing Ca^2+^/Mg^2+^). The cells were subsequently resuspended in 100 µl of FACS buffer with a secondary anti-mouse-APC antibody (BioLegend, catalogue number-405308). The cells were incubated at 4 °C for 30 minutes on ice and washed with PBS as before. 10 µl was taken from each sample for RBC analysis and the rest of the sample was fixed with 4% paraformaldehyde (PFA) and incubated on ice for 20 min. The sample was then washed with PBS and subsequently treated with ACK lysis buffer (Gibco, catalogue number-A10492-01) to perform analysis of RBCs. The sample was washed and resuspended in 500 µl of FACS buffer. In the forward and side scatter profile, monocytes and neutrophils were gated for the analysis. For gating of monocytes forward and side scatter pattern was used. The surface markers were not used for this gating.

### Glycan array analysis for the binding of HMGB1 with sialic acids

Chemoenzymatically synthesized sialyl glycans were quantitated utilizing DMB-HPLC analysis and were dissolved in 300 mM sodium phosphate buffer (pH 8.4) to a final concentration of 100 µM. ArrayIt SpotBot^®^ Extreme was used for printing the sialoglycans on NHS-functionalized glass slides (PolyAn 3D-NHS slides from Automate Scientific; catalogue number-PO-10400401). Purified mouse anti-HMGB1 antibody (BioLegend; catalogue number-651402, Lot# B219634) and Cy3-conjugated goat anti-mouse IgG (Jackson ImmunoResearch; catalogue number-115-165-008) were used. Fresh HEPES buffer (20mM HEPES, 150mM NaCl ± 500µM ZnCl_2_) was prepared immediately before starting the microarray experiments.

Method described in (34) was adapted to perform the microarray experiment. Each glycan was printed in quadruplets. The temperature (20 °C) and humidity (70%) inside the ArrayIt^®^ printing chamber was rigorously maintained during the printing process. The slides were left for drying for an additional 8 h. Printed glycan microarray slides were blocked with pre-warmed 0.05 M ethanolamine solution (in 0.1 M Tris-HCl, pH 9.0), washed with warm Milli-Q water, dried, and then fitted in a multi-well microarray hybridization cassette (ArrayIt, CA) to divide it into 8 subarrays. Each subarray well was treated with 400 µl of ovalbumin (1% w/v) dissolved in freshly prepared HEPES blocking buffer ± 500 µM of Zn^2+^ (pH adjusted for individual experiments) for 1h at ambient temperature in a humid chamber with gentle shaking. Subsequently, the blocking solution was discarded, and a solution of HMGB1 (40 µg/ml) in the same HEPES buffer (± Zn^2+^, defined pH) was added to the subarray. After incubating for 2 hours at room temperature with gentle shaking, the slides were extensively washed (first with PBS buffer with 0.1%Tween20 and then with only PBS, pH 7.4) to remove any non-specific binding. The subarray was further treated with a 1:500 dilution (in PBS) of Cy3-conjugated goat anti-mouse IgG (Fc specific) secondary antibody and then gently shaken for 1 hour in the dark, humid chamber followed by the same washing cycle described earlier. The developed glycan microarray slides were then dried and scanned with a Genepix 4000B (Molecular Devices Corp., Union City, CA) microarray scanner (at 532 nm). Data analysis was performed using the Genepix Pro 7.3 analysis software (Molecular Devices Corp., Union City, CA).

### Purification of HMGB1 from *E. coli* and HEK293 freestyle

Expression and purification of full-length murine His-HMGB1 in *E. coli* were performed as described before (35). Mutagenesis was performed using a QuikChange site-directed mutagenesis kit (Agilent).

For HMGB1 expression in mammalian cells, the complete open reading frame of murine HMGB1 was cloned into pcDNA3.1(+)-C-6His vector (GenScript). Transfection was performed using FectoPRO transfection reagent (Polyplus transfection). Recombinant His-HMGB1 was produced in 293-freestyle cells (ThermoFisher Scientific). Purification of His-HMGB1 from 293-freestyle cell lysate was carried out using Ni SepharoseTM 6 Fast Flow gel (GE Healthcare). After purification, His-HMGB1 was 99% pure as judged by silver staining.

## Acknowledgments

We thank Sandra Diaz and Patrick Secrest for their excellent technical help with the work.

## Author contributions

SS Siddiqui, C Dhar, V Sundaramurthy and A Varki wrote the manuscript. SS Siddiqui, C Dhar, V Sundaramurthy, A Sasmal, H Yu, X Chen, E Bandala-Sanchez, LC Harrison, X Zhang, M Li and D Xu designed and performed the experiments. All authors reviewed the manuscript and approved it.

## Supplementary data

**Supplementary Fig 1:**
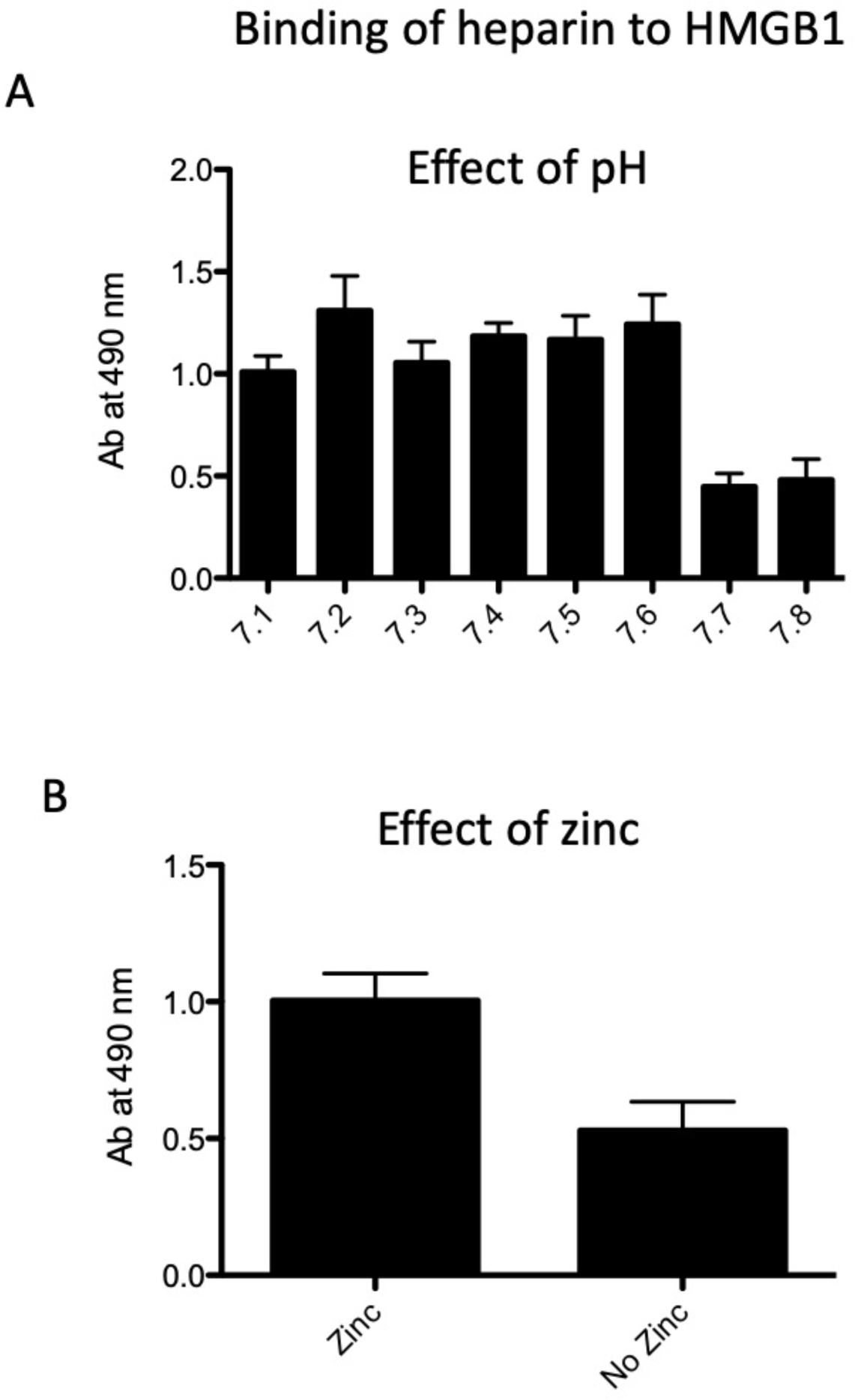
Previously known binding partner of HMGB1, another anionic glycan heparin does not exhibit this extreme pH sensitivity, and zinc only partially facilitates binding: A) The binding of HMGB1 with heparin was determined by ELISA using a binding buffer at different pH ranges (7.1-7.8) B) The binding assay of HMGB1 and heparin was also performed with a binding buffer with and without zinc. The experiment was performed in triplicate, where data shows mean±SD.

**Supplementary Fig 2:**
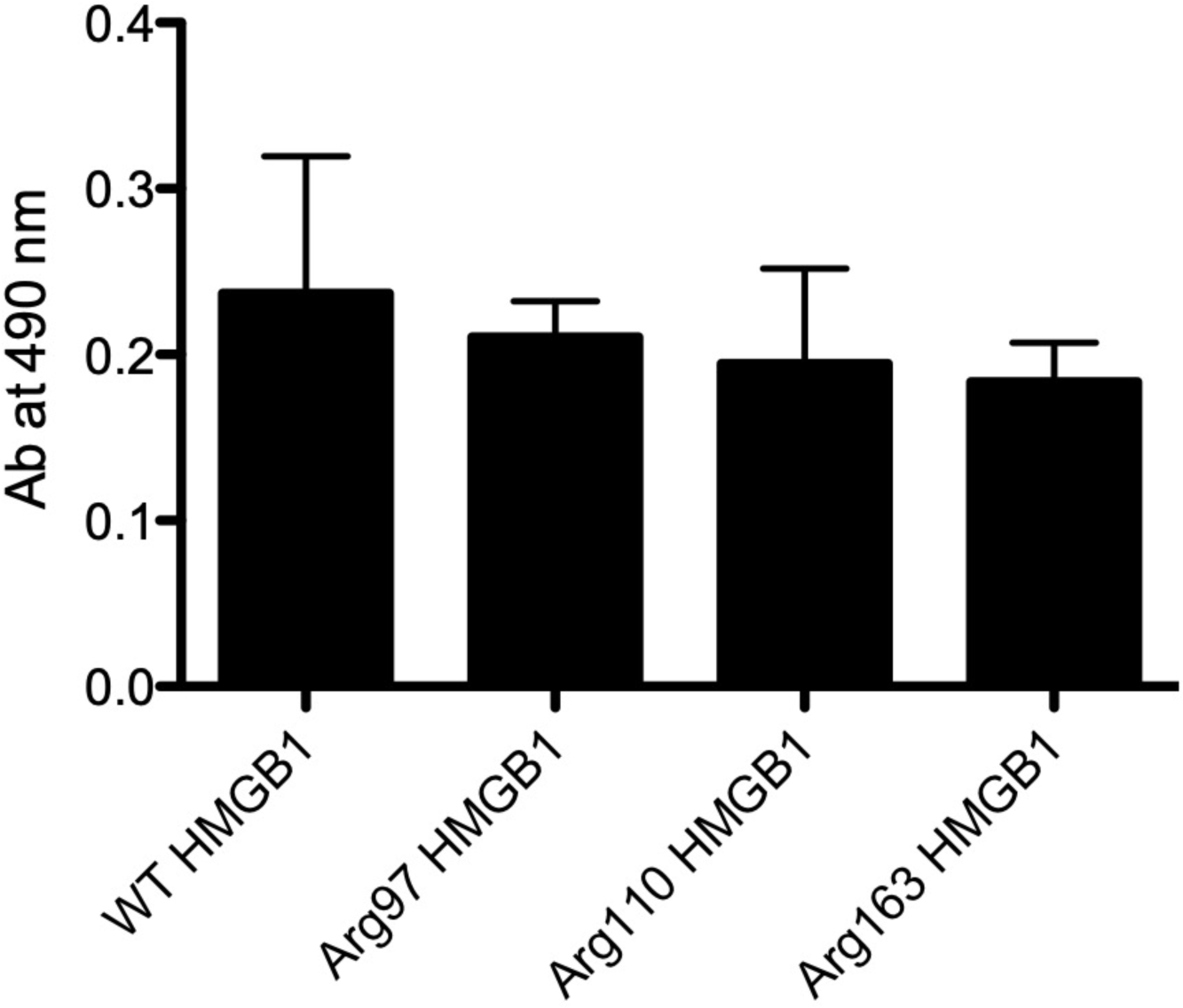
Binding of sialic acid with different arginine mutants of HMGB1: Different arginine mutants of HMGB1 B-box were generated and binding of these mutants with 3’-sialyllactose was measured by ELISA. The experiment was performed in triplicate, where data shows mean±SD.

**Supplementary Fig 3:**
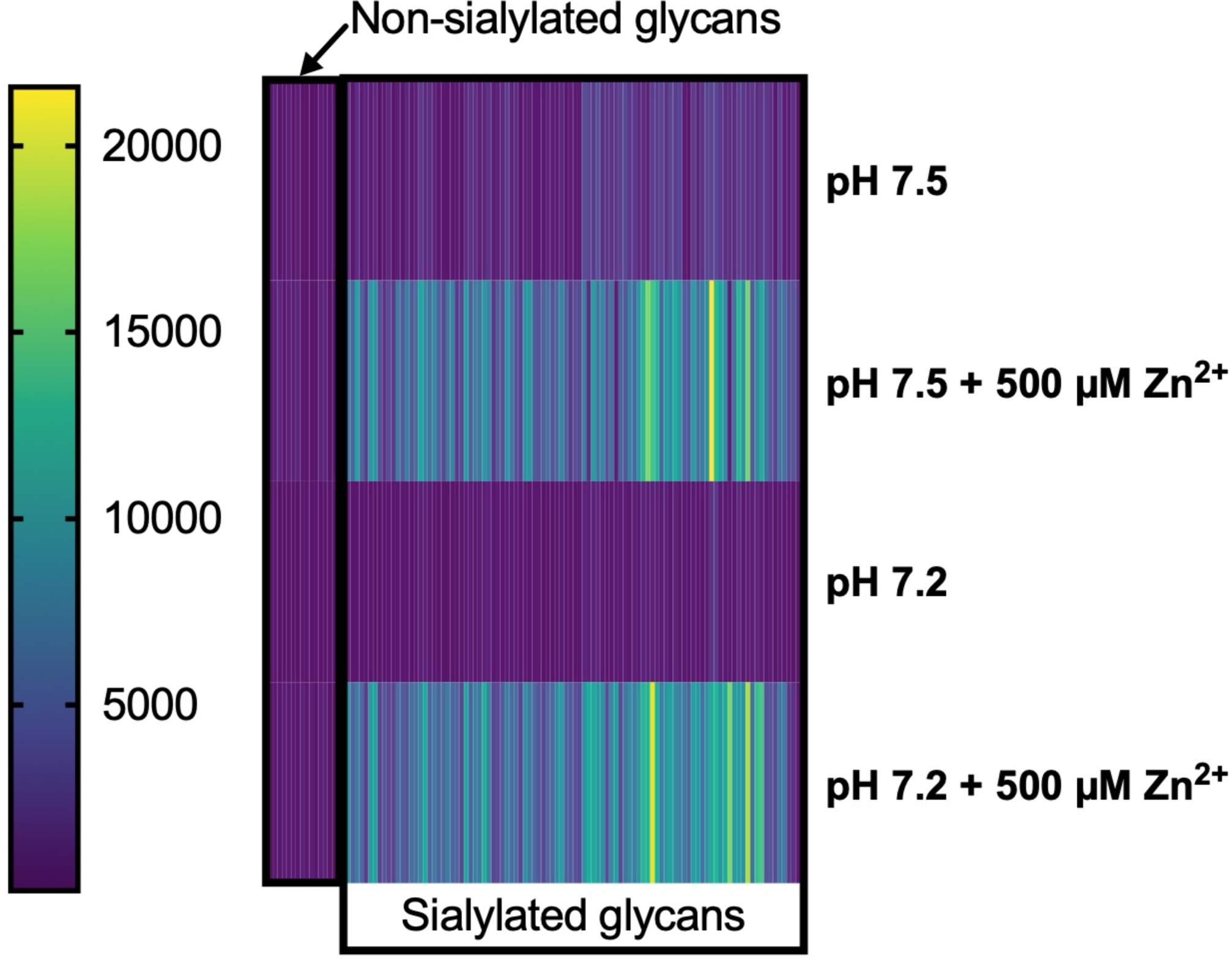
Heatmap representing average RFUs of HMGB1 binding to sialosides and non-sialosides under various conditions.

**Supplementary Fig 4:**
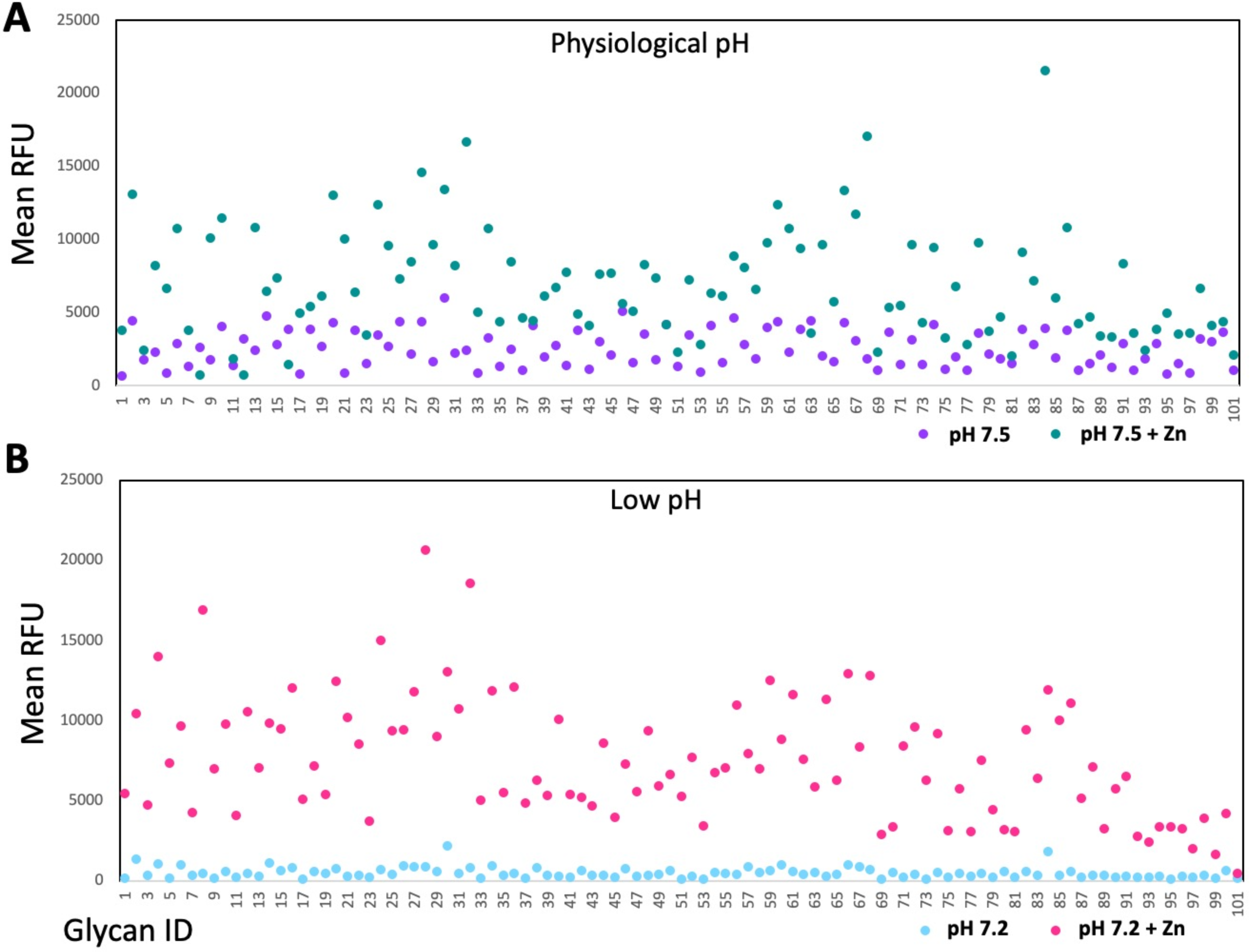
Binding of HMGB1 with sialic acids at physiological zinc concentration: The binding of HMGB1 with sialoglycan probes on a glycan array was performed using different zinc concentrations. The data shows mean RFU ± SD.

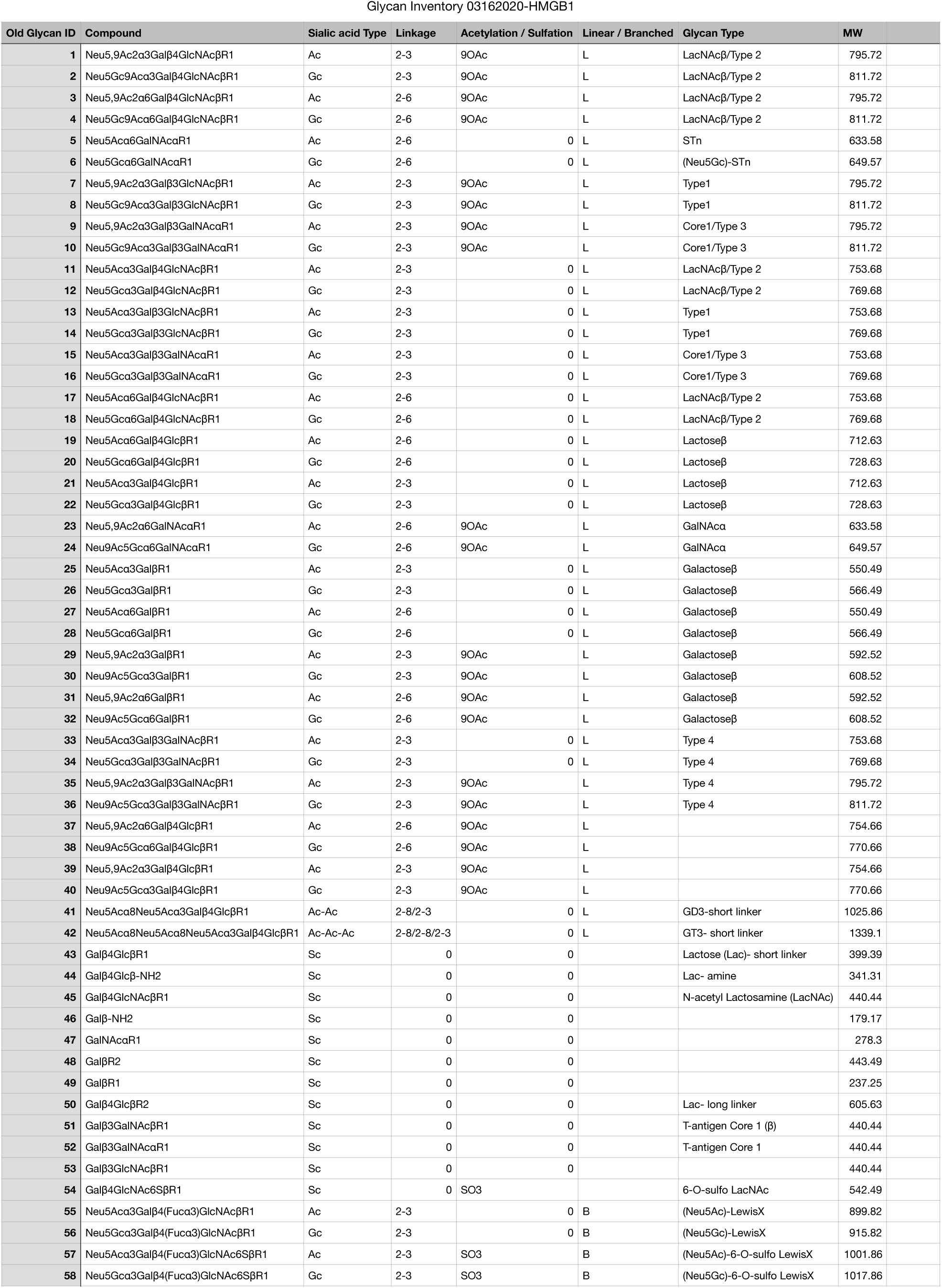
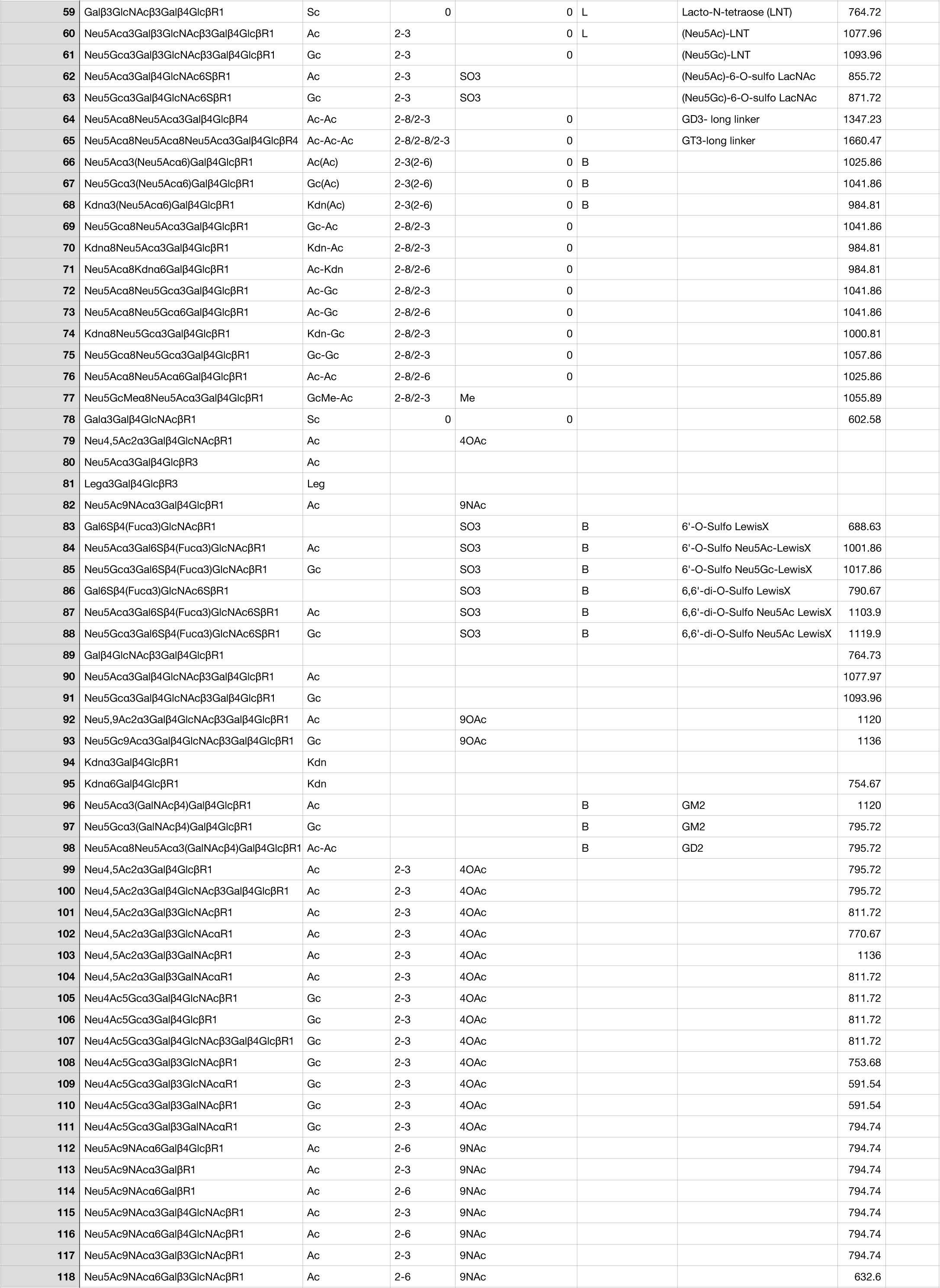
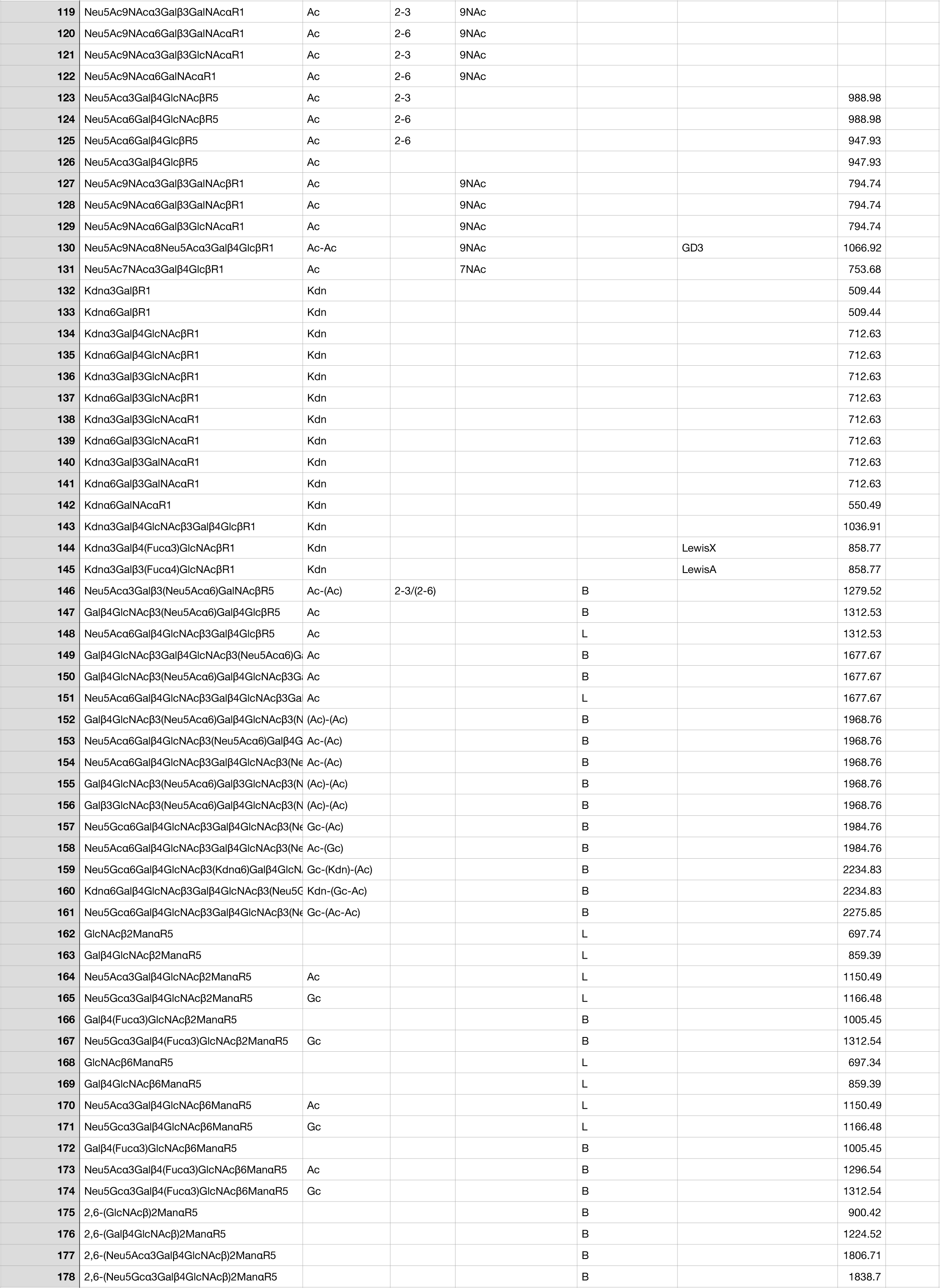
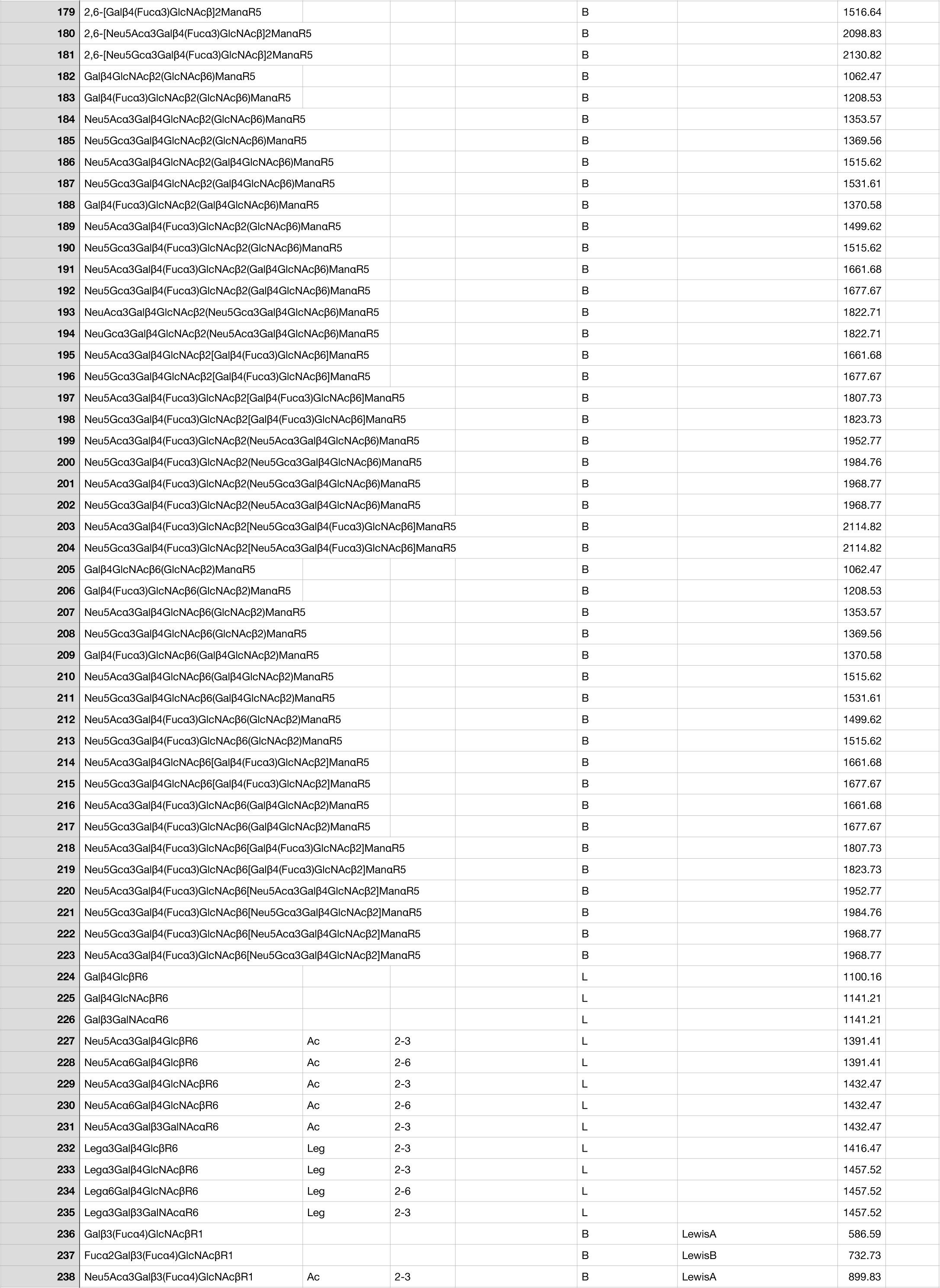
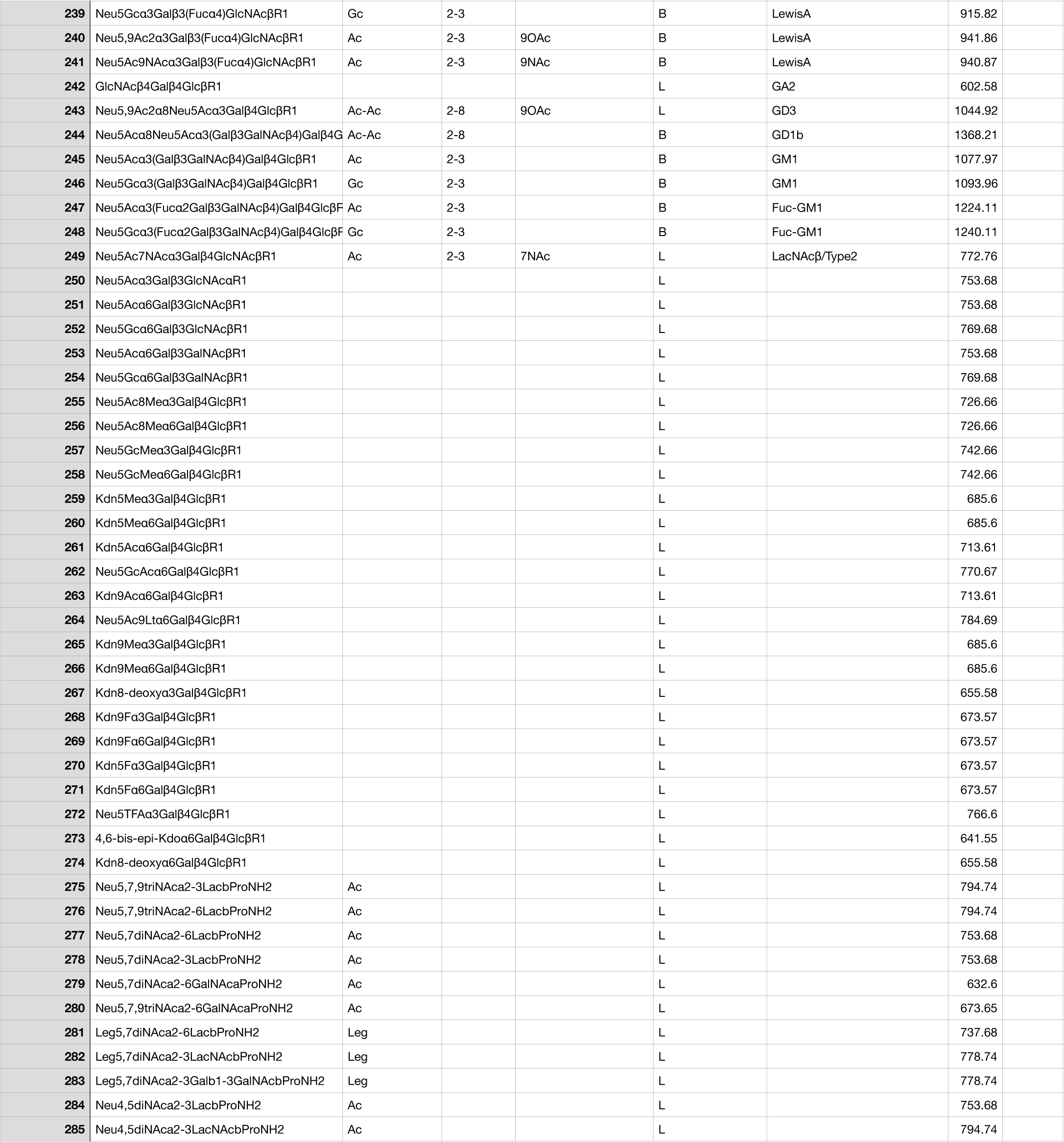

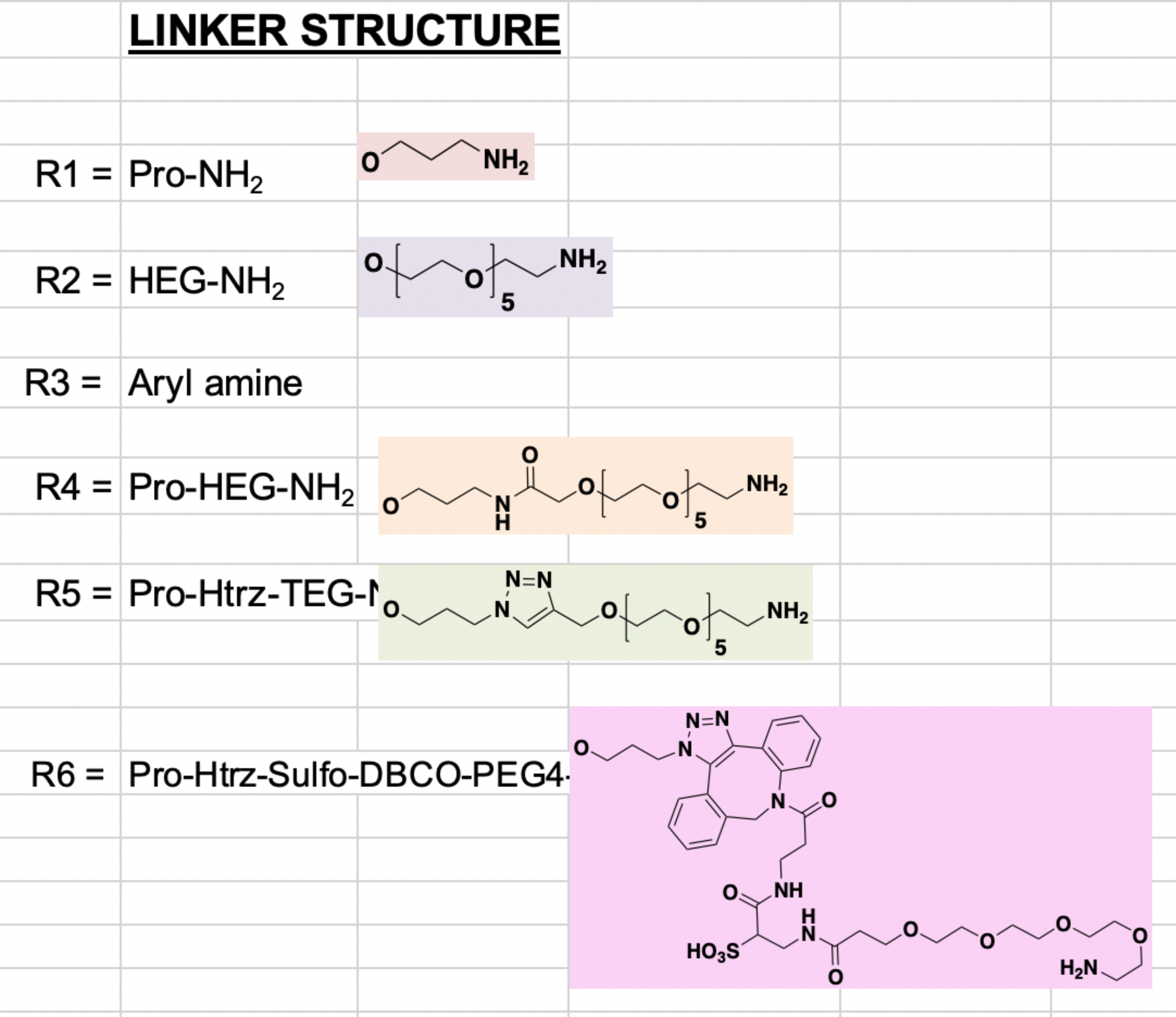

